# Aging increases lateral but not local inhibition of orientation processing in primary visual cortex

**DOI:** 10.1101/067264

**Authors:** Zhengchun Wang, Shan Yu, Yu Fu, Yifeng Zhou, Tzvetomir Tzvetanov

## Abstract

Aging-related declines in vision can decrease well-being of the elder. Concerning early sensory changes as in the primary visual cortex, physiological and behavioral reports seem contradictory. Neurophysiological studies on orientation tuning properties suggested that neuronal changes might come from decreased cortical local inhibition. However, behavioral results either showed no clear deficits in orientation processing in the elder, or proposed stronger surround suppression. Through psychophysical experiments conducted on old and young human subjects combined with computational modeling, we resolved these discrepancies by demonstrating stronger lateral inhibition in the elder while neuronal orientation tuning widths, related to local inhibition, stayed globally intact across age. We confirmed this later finding by re-analyzing published neurophysiological data from rhesus monkeys, which showed no systematic tuning width changes, but instead a higher neuronal noise with aging. These results suggest a stronger lateral inhibition and mixed effects on local inhibition during aging, revealing a more complex picture of age-related effects in the central visual system than previously thought.

**Significance Statement:** Visual functions decline during aging, adversely affecting quality of life. Much of this dysfunction is probably mediated by disturbances in the balance between inhibition and excitation in the central visual system. It was proposed that the inhibitory function within the aging visual cortex might be modified, but huge discrepancies exist among different reports. Here we identify the specific inhibitory circuit change, which has not been clearly evaluated, by using behavioral measures, neural modeling and re-analysis of non-human primate electrophysiological data. We provide evidence that a stronger lateral inhibition and mixed effects on local inhibition during aging, revealing a more complex picture of age-related effects in the central visual system than previously thought.

**Author contributions:** ZCW and TT designed and analyzed behavioral experiments; ZCW performed experiments; ZCW and TT performed data analysis and neurophysiological re-analysis; YS and FY provided physiological data and discussed its re-analysis; TT performed modeling; ZCW, YFZ and TT provided project supervision and funds; ZCW and TT wrote the paper; all authors discussed and commented on the manuscript.

## Introduction

Human cognitive abilities decline during aging (Hedden and Gabrieli, 2004; Craik and Bialystok, 2006; Mather, 2010), but also early sensory processing is affected. Age-related modifications in early neuronal processing and perceptual abilities of visual system have been intensively studied in the last 25 years (Owsley, 2011; Andersen, 2012). However, there is no clear unifying account of early stages changes, especially specific inhibitory circuit change, as in spatial vision.

Neurophysiological studies suggested that a reduction in inhibitory function might be related to age-related perceptual dysfunctions in spatial vision (Schmolesky et al., 2000; Leventhal et al., 2003). Much of the inference was drawn from observations in animal research on neuronal orientation response properties of classical receptive field (CRF) (Somers et al., 1995; Shapley et al., 2003), and there seems to be a broad agreement for altered cellular properties that are known to depend on inhibition. Nevertheless, some behavioral studies showed that there are low-level processes of visual functions which are inconsistent with the physiological interpretation of decreased inhibition: (1) psychophysical sensitivities to orientation processing in elder populations are not systematically and substantially changed when compared to younger adults (Betts et al., 2007; Delahunt et al., 2008; Govenlock et al., 2009); (2) surround suppression in the elder was found increased, which was interpreted with stronger inhibitory interactions (Karas and McKendrick, 2015; Nguyen and McKendrick, 2016). It appears that there are two levels of early visual processing in spatial vision that need to be simultaneously accounted for: (i) a local inhibition related to CRF and (ii) a form of lateral inhibition related to long-range interactions among neurons (Gilbert, 1992; Spillmann and Werner, 1996; Angelucci et al., 2002; Angelucci and Bressloff, 2006).

Interestingly, these two levels of interactions (Angelucci and Bressloff, 2006) can be accessed non-invasively through the use of psychophysical measures and computational modeling (Gilbert and Wiesel, 1990a; Tzvetanov and Womelsdorf, 2008; Tzvetanov, 2012). We used the tilt illusion effect where the presence of an orientated surround stimulus biases the perceived orientation of a simultaneously presented center test. Current opinions propose that it is explained through lateral inhibition between spatially arranged orientation hypercolumns of neurons in V1 (Georgeson, 1973; Wenderoth and Smith, 1999; Kapadia et al., 2000) (Figure 1a). Specifically, in the classical center-surround tilt repulsion effect (Gilbert and Wiesel, 1990a) (Figure 1a), one can infer the local interactions through the indirect measure of the orientation tuning widths of a theoretical population of neurons (Figure 1a, and 1b, top), which is directly related to the local inhibition within one hypercolumn (Somers et al., 1995), but also the lateral interactions between different hypercolumns (blue arrows in Figure 1a). Both local and lateral interactions affect the tilt repulsion curve (Figure 1b, bottom), that describes the subject’s misperception of a vertically oriented center target when the orientation of the surround is varied (Figure 1a, gray stimulus). Through model adjustment to the behavioral data, estimated mean values of these physiological variables of the neural population was accessed for each person.

**Figure 1:**
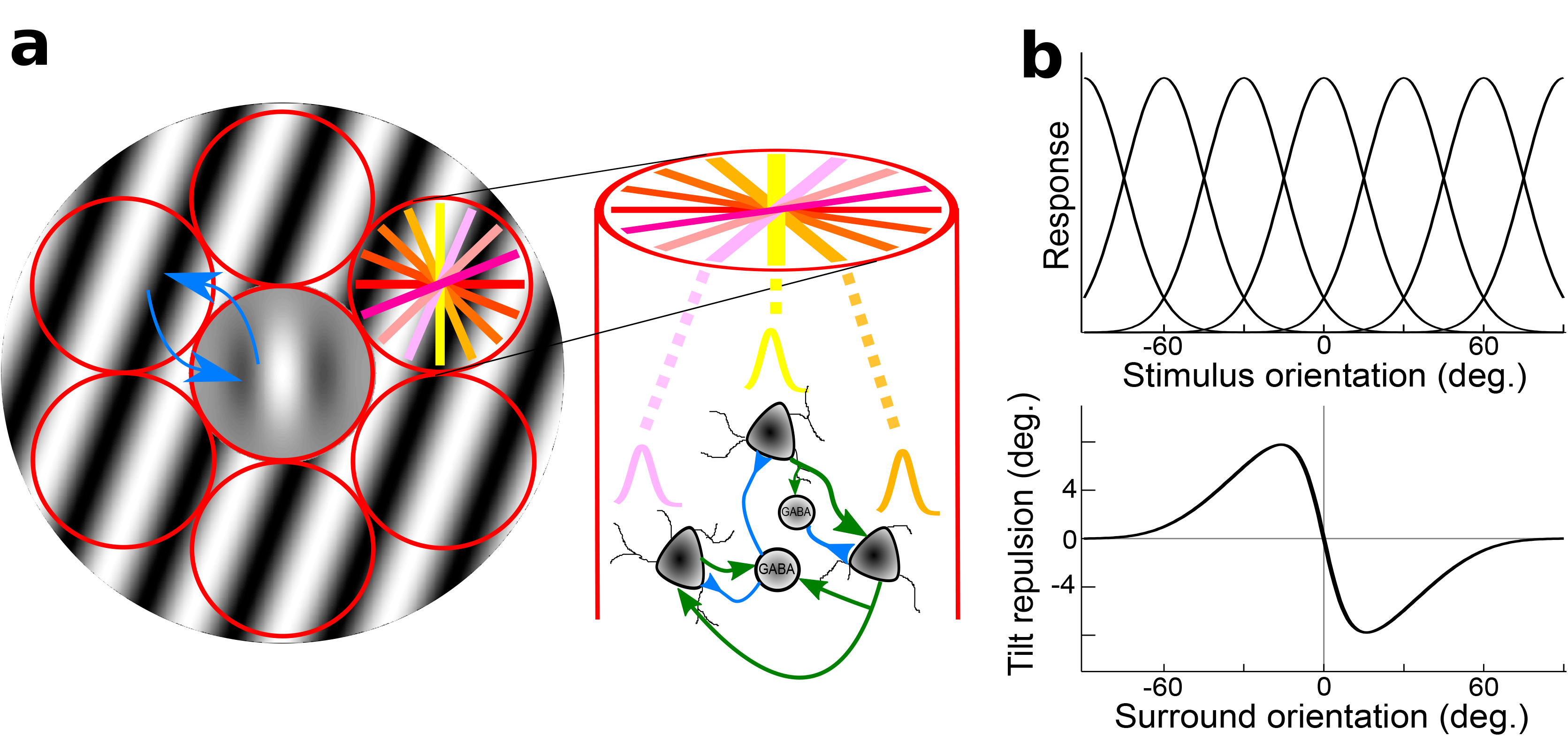
Illustration of center-surround stimulus, orientation hypercolumns, population tuning curves and behavioral outcome. (a) Illustration of the stimulus, a center Gabor patch surrounded by an annulus of oriented grating, padded with orientation hypercolumns (red circles); the colored lines in right-top circle depict different preferred orientations of the local neuronal population; on the right, illustration of local interaction within an orientation hypercolumn, with three example neurons with different preferred orientations. Two inter-neurons, and their local connections (blue/green are inhibitory/excitatory connections); blue arrows depict inhibitory lateral interactions between hypercolumns. (b) Theoretical population orientation tuning curves (top) together with typical tilt repulsion curve (bottom) describing the orientation misperception of a vertical center stimulus as a function of the orientation of the surround; zero is vertical and positive values are clockwise tilts.

Here, we carried psychophysical measures in young and elder populations in order to characterize their center-surround tilt repulsion at two spatial frequencies (SFs), obtained from the individual contrast sensitivity function (CSF). Then, we performed modeling of subjects’ perception assuming it is obtained from decoding these V1 neuronal activities. Contrary to the common neurophysiological belief that age-related changes of visual perception in the elder mainly stem from a reduction in inhibitory functions, the combination of psychophysical measures and modeling showed different aging-related effects on different kinds of inhibition: (1) an increased lateral inhibition with aging and (2) maintained local orientation tuning widths, and thus local inhibition, of the neuronal population in the elder, which can be investigated directly in electrophysiological studies. We, then, re-analyzed previously published neurophysiological data about orientation processing from our laboratory and extracted the tuning width of neurons from young and old macaques. These physiological results were consistent with our behavioral and computational findings of unchanged orientation tuning widths.

## Materials and Methods

### Subjects

The current study included two groups: 20 (7 females) younger adults at age of 22 to 40 (26.45 ±3.73) years and 20 (7 females) older adults at age of 65 to 77 (69.55 ± 3.63) years. One young subject data were excluded due to no clear staircase convergence in the tilt illusion task (very high thresholds). One elder’s data were excluded because of incorrect task performance in the replication of the contrast discrimination task (staircases without convergence properties). Subjects younger than 30 years old were students of the University of Science and Technology of China, while others were recruited from local communities. All subjects were naive to the purpose of the experiments (except the author), and their informed consent was obtained before participation. Examination (MMSE) was performed on elder subjects to exclude probable dementia. Alcoholism, stroke and depression were also exclusion criteria by questionnaires before conducting experiments. Participants also provided information about their general health, to exclude people with systemic conditions known to affect visual function (for example, diabetes, migraine, schizophrenia, and epilepsy) or who were taking medications known to affect visual function (e.g., anti-anxiety or anti-depressant medications). All participants were measured with normal or corrected-to-normal visual acuity (younger=1.05±0.12, elder=1.05±0.17 (MAR); mean±std). This research has been approved by the ethics committee of University of Science and Technology of China and followed the guidelines of the Declaration of Helsinki. Subjects were paid for their participation on an hour basis.

### Set-up

The experiment was conducted in a dimly illuminated room. Stimuli were displayed on a 40.0 cm×30.0 cm CRT monitor (Sony G520; 85 Hz, resolution of 1600×1200 pixels) with self-programmed Matlab functions (Mathworks Inc.) using the Psychophysics toolbox (Brainard, 1997). Stimuli were displayed using an NVIDIA Quadro K600 graphics system and viewed binocularly. The screen area was delimited by a circular window of diameter 30.0 cm cut in a black cardboard centered on the screen in order to avoid local cues for vertical/horizontal and position (Tzvetanov, 2012). Luminance values were obtained with the help of the contrast box switcher (Li et al., 2003), that allowed to extend luminance range digitization above 10 bits, and thus provided the necessary minimum contrast step for contrast detection measurement. Calibration was performed each day of measurement throughout the experiments.

The eye-to-screen distance was maintained using a chin rest and fixed at 4 meters for CSF measurement and the experiments for tilt illusion test with high-SF, and 2 m for tilt illusion test at low-SF and the replication study of contrast discrimination.

### Stimuli

The stimuli used for CSF measurements were vertical sine-wave gratings with different SFs (0.71, 1, 1.41, 2, 2.83, 4, 5.66, 8, 11.31, 16 and 22.63 cycle/degree) for both groups (Figure 2a). These stimuli subtended 2° aperture, and were presented on a mean luminance of 40 cd/m^2^ background. To minimize edge effects, a border-mask was used to blend the stimuli to the background. From each CSF, a low and a high SFs were chosen for the following tilt illusion measures.

**Figure 2:**
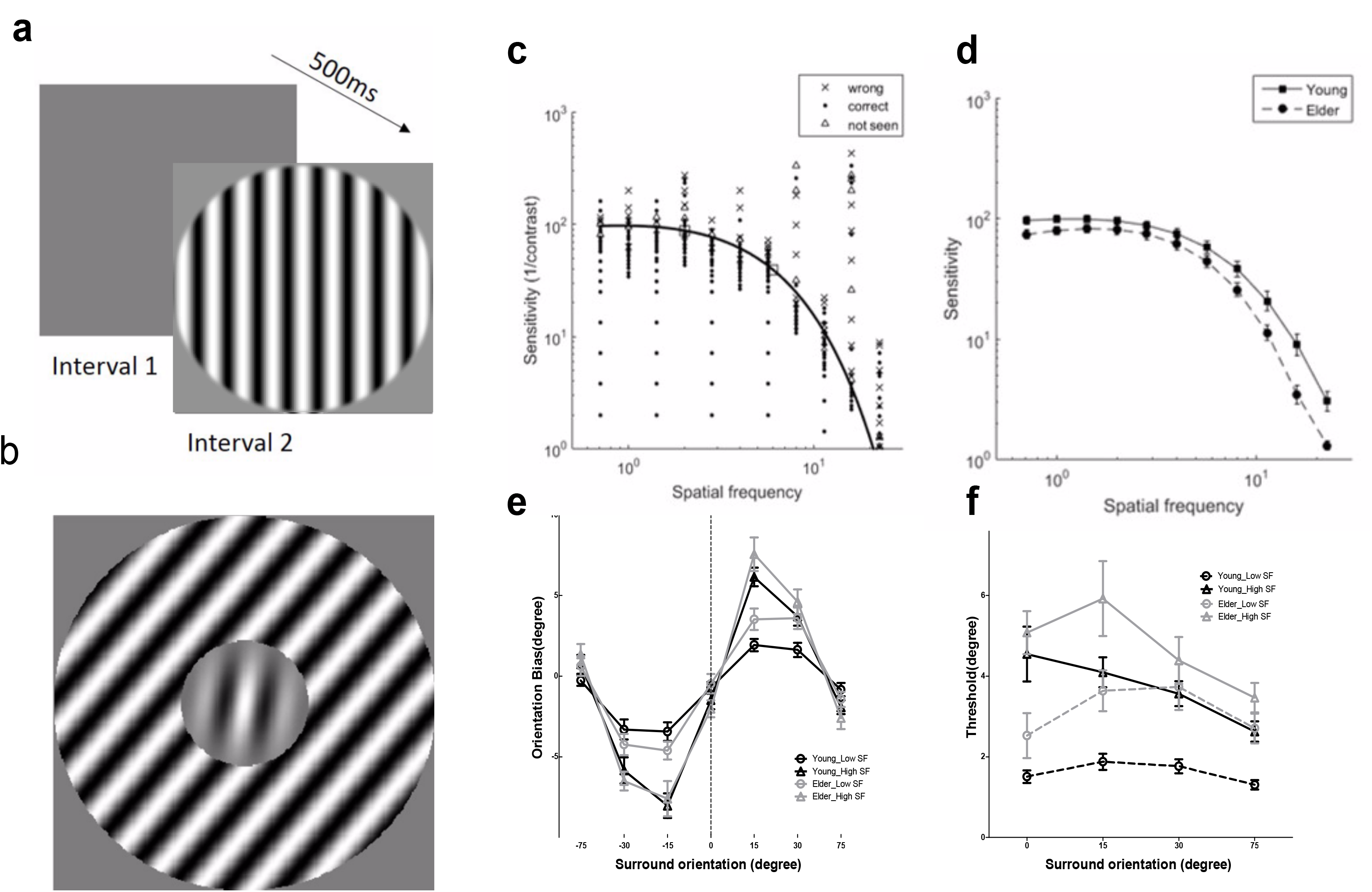
Examples of stimuli used and results of contrast sensitivity function and tilt illusion measures. (a) Schematic illustration of the contrast sensitivity function (CSF) measure; (b) Example of stimuli in tilt illusion measure. (c) Examples of results of CSF measures and fits in the 2-Alternative unforced Choice (2AuFC) design; squares depict the two chosen SFs for the subsequent tilt measures; (d) Averaged values of contrast sensitivities at the eleven different SFs measured for the elder and younger groups; (e) Tilt illusion results, indicated by orientation bias necessary to perceive the center as vertical, as a function of SOs and SFs (low-SF: circles; high-SF: triangles) for the elder (gray) and younger (black) groups; negative (positive) deviations for negative (positive) surround orientations correspond to repulsion; (f) Orientation thresholds around perceived verticality.

The stimuli used in the tilt illusion experiments were a central Gabor patch (test grating) surrounded by an annulus of the inducing sine-wave grating with different orientation (0°, ±15°, ±30°, ±75°; angle is defined with respect to the orientation of the center) (Figure 2b). For both center and surround, stimuli had same SF and contrast (90%) and were presented on a mean background luminance of 35 cd/m^2^. Two spatial frequency values (low-SF and high-SF), obtained from the CSF, were used as spatial frequencies for the tilt illusion measurement. The size of the stimuli were scaled with SF, keeping the central window diameter of the target stimulus fixed at 4 cycles. Surround annulus width was equal to center diameter. For both center and surround, the cosine had a phase of zero. The orientation of the central Gabor patch was varied around vertical from trial to trial to measure subject’s perceived vertically. The Gabor patch was defined through the following equation:

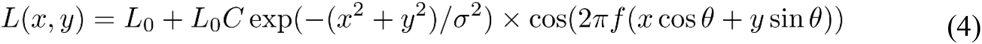

with *L*_0_ the background luminance of the screen, *C* the Gabor patch contrast, and *f* and *θ* its spatial frequency and angle relative to vertical.

The stimuli used in the center-surround contrast discrimination experiments consisted of a small central Gabor patch of vertical orientation (f=4 c/deg, equ.4), with its diameter fixed at 4 cycles. It was presented either alone or surrounded by an annulus of sine grating (4 c/deg, inner radius equals center Gabor patch radius, outer radius is inner radius plus center diameter). All phases were fixed at zero. Stimuli were presented on a mean background luminance of 35 cd/m2. The stimulus with center alone was a test Gabor patch whose contrast was varied in order to measure the perceived contrast of the center when it is flanked by the surround. The center-surround stimulus had a predefined set of possible contrasts. The full measurements included all combinations of contrasts for center-surround among the three values of 20%, 40% and 80%. This was performed in order to compare our design to a recent study (Karas and McKendrick, 2015), but for the present report we restrict our presentation to the center-surround of 80%−80% that matches closely the tilt illusion condition.

### Procedure

Subjects were instructed before each experiment of the currently requested task and to keep fixation on the center dot during stimulus presentation. Before measurement, each subject received a short practice session (about 80 trials). Experiments were initiated by subjects with a keyboard press and stimuli were presented 200 ms after fixation point disappearance. Subjects responded by pressing corresponding keyboard keys. A CSF measurement was conducted prior to tilt illusion measurement. Tilt illusion measurement for high- and low-SF were conducted in two separate blocks. A center-surround contrast discrimination task was also conducted for consideration of comparisons to previous reports.

For CSF measurement, there were 275 trials in total (25 trials for each SF value, total 11 SFs). Each trial consisted of two intervals separated by a 500 ms inter-stimulus interval (ISI), and each interval was announced by a short beep. The target vertical sine-wave grating was presented in one of the two intervals for 100 ms. A two-alternative-unforced-choice (2AuFC) design with 3 response keys was used in CSF measurement (Garcia-Perez, 2010). The observer was required to indicate in which interval appeared the grating by pressing two corresponding keys, or a third key for ‘not seen’ if the subject didn’t know in which interval the grating appeared. For the staircase procedure, these key presses were randomly drawn as correct/incorrect responses. Feedback was provided by different sound for correct and incorrect responses and mute for the undecided key. Psychometric curves were measured using the weighted up-down adaptive procedure (Kaernbach, 1991b) with steps up/down of 6.4/0.8 for SFs of 1 to 16 c/d in one octave steps and 6/2 otherwise (base contrast step change was 10%). Starting points were contrasts of 0.5, 0.005, 0.5, 0.005, 0.5, 0.005, 0.5, 0.005, 0.7 0.05, 0.8 for the successive 11 SFs, respectively, and each staircase “down” step-size was additionally 3 times bigger for the first 4 trials (example results in Figure 2c).

For tilt illusion measurement, there were two blocks and each block consisted of 420 trials (60 trials for each SO, total 7 SOs). The stimulus in each trial was presented for 9 frames and no feedback was provided. SFs used in the tilt illusion measurement were chosen from subject’s CSF outcomes: one near peak CSF, called low-SF, and one higher, called high-SF, with the condition that it was not too close to the cut-off SF, above which the subject cannot see full contrast stimuli. The 3 keys design was used where the observer was required to report the orientation (clockwise/counter clockwise) of the center test Gabor patch from his/her internal vertical standard by pressing two predefined keys. The third key was allowed if the subject felt he/she could not perceive any orientation in the central part (due to surround suppression; especially at high SFs), and “not seen” cases were randomly drawn as clockwise/counter clockwise for the staircase procedure. The center orientation was varied according to the weighted up-down rule with steps up/down of 5/2 and 2/5 and base step of 1 degree.

The procedure for contrast discrimination task was as follows: one block consisted of 300 trials measuring the three psychometric functions for the center-surround stimulus with three reference center contrasts and surround contrast fixed within the block. In each trial, there were two intervals, separated by 500 ms, presenting in one interval the central grating alone and in the other the central grating surrounded by an annulus. The stimuli were randomly assigned to intervals. Each stimulus was accompanied by a short beep and presented for 100 ms. No feedback was provided. The test contrast was varied with the weighted up-down staircase procedure (Kaernbach, 1991a) and for each reference contrast four staircases were ran with steps up/down of 7/1.5, 1.5/7, 6/2.5, and 2.5/6 of the baseline step of 5% contrast change. Starting points were 0.05, 0.9, 0.05 and 0.9 respectively. The 2AuFC task with three key responses was used where the observer was required to nominate in which interval (first/second) the central patch had the highest contrast. The third key was used by observers if they could not identify in which interval the grating had higher contrast and “indecision” cases were randomly drawn as the test perceived lower/higher contrast.

### Data analysis

We used Bayesian fitting to adjust theoretical psychometric functions to the data (Treutwein and Strasburger, 1999).

#### Contrast sensitivity function

A 2D psychometric function was fit to the 2D contrast-SF (*c, f*) data, with the probability of correct response defined as:

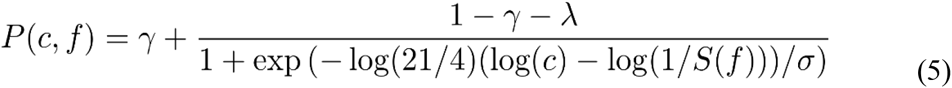

with parameters y and λ being subject’s “guess rate” (see below) and lapsing rate, and 2σ defining the spread between 16%−84% of the function in the range γ to 1-λ (assuming constant spread, σ, across SFs). *S*(*f*) is the standard 3-parameters sensitivity function (Rohaly and Owsley, 1993):

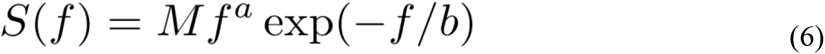

used in previous studies to define the CSF shape in the SF dimension. The 3 response keys design data was processed in the “Fechner paradigm”, with the 3rd key presses considered as half-correct and half-wrong (Garcia-Perez and Alcala-Quintana, 2011), which in the event the subject follows the 3rd key instructions allows a decrease in measurement variability. The Likelihood function is then:

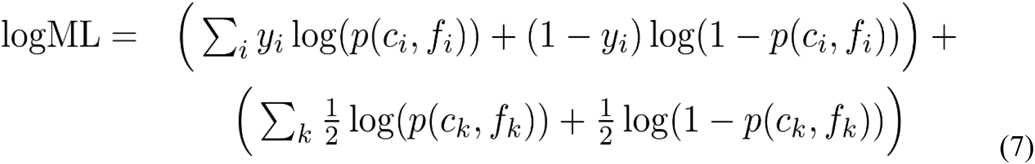

with the first sum (trials with index *i*) running over all responses 1st or 2nd interval and the second sum (trials with index k) running over the “undecided” 3rd key responses. The variables in the above equation are: (*c_i_*, *k, f_i,k_*) – contrast & SF pair presented at trial *i* or *k, y_i_* – correct/incorrect (1/0) response of subject at contrast & SF levels (*c_i_, f_i_*), *p* (*c_i_, k, f_i_ k*) – theoretical probability of correct response (psychometric function). In this ML equation, the first term is the standard log-ML term for fitting binomial data; the second “Fechner” term is simply log(*p*(1–*p*)), the logarithm of the binomial variability at the stimulus levels for which the subject pressed “undecided”/ “not seen”; it is maximized when p(*c_k_, f_k_*) is 0.5, i.e., when the subject is totally uncertain about the interval of signal presentation, and thus provides a firm theoretical ground for introducing “undecided” / “not seen” responses into the 2AFC design. The lapsing rate was fixed at 1% for all but one subject where it was zero. The “guess rate” γ was 0.5 and an example CSF fit for one subject is displayed on Figure 2c, that also displays the responses given by the subject to each presented stimulus.

#### Tilt repulsion

We fit a 1D psychometric function to the orientation discrimination data for each surround orientation, with probability of CW responses to target orientation *θ* given by:

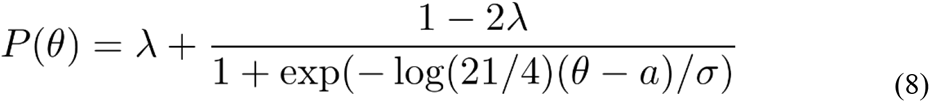

where λ is subject’s lapsing rate, and *a* and *σ* being the perceived vertical orientation (also called “bias”) for the given surround and the threshold of the subject for perceiving a deviation from verticality, respectively. Because of the symmetry in the experimental design (symmetric surrounds of ±15, ±30, ±75 degrees), for the fitting we imposed that thresholds of opposite surround orientations (e.g. −30 and +30 degrees) are the same. The lapsing rate was fixed at 0% for low-SF condition, and 1% for high-SF. The data was processed by eliminating any datum with 3rd key responses (subject did not see the target), and we computed the amount of surround suppression as the proportion of 3rd key presses. Bias values were computed as the half-difference between two opposite surround orientations.

#### Center-surround contrast perception

The 80%–80% condition was fit with a logistic psychometric function:

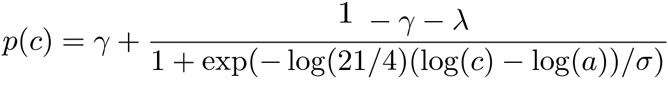

representing the probability to respond test Gabor patch (without surround) had higher contrast. We used the “Fechner’s” definition for undecided key presses into the maximum-likelihood equation (“undecided” cases were considered as half one category and half the other, e.g. (Garcia-Perez and Alcala-Quintana, 2011) Garcia-Perez & Alcala-Quintana, 2011, equ.7). The parameters *γ* and *λ* were constrained as follows: *λ* was constrained between 0 and 0.5, while *λ* was constrained between 0 and *λ_max_* (with *λ_max_*=1-*γ*-(1–2*λ*)/2/(1+exp(log(21/4)log(*a*)/*σ*)) in order to have *p*(*c*=1)≥0.5), and both had flat prior. The second constrain was necessary because we found that multiple subjects (16/40) had difficulties to discriminate the two central patches for test targets (no surround) of contrasts around 90% (near the maximum available), and disambiguate parameters entanglement (that is, their psychometric functions were not saturating near c=1). A suppression ratio was calculated (perceived contrast/physical reference contrast) to quantify the strength of the center-surround interactions (Karas and McKendrick, 2015). A value below 1 indicates surround suppression, while a value above 1 indicates enhancement.

### Statistical Analysis

For contrast sensitivity, a two factors between-within ANOVA (age group × spatial frequency) was conducted. For tilt illusion, a 3-way between-within subject ANOVA (age group × surround orientation × spatial frequency) was conducted on all data. For center-surround contrast discrimination task, a t-test was used to analyze the suppression ratio data (paired or unpaired, as appropriate). All statistical levels use Geisser-Greenhouse epsilon-hat adjusted values, where appropriate. Spearman rank correlation was used. Data are expressed as mean±SE.

### Model

The model was developed for a previous study in our laboratory, concerning Amblyopia deficits (Tzvetanov, Huang, Liu, Liu, Zhou, Chinese Neuroscience Society 11th biennial meeting, WuZhen, 2015), and a detailed description was given in the accompanying manuscript (Huang, Zhou, Wang, Wang, Liu, Liu, Tzvetanov, submitted). We provide the model description as in the original work for consistent methodological description in the current work.

#### Simple model of V1 surround-to-center interactions

We assume, as in many previous studies, that simple feature perception as local orientation and contrast can be explained through the decoding of primary visual cortex neuronal activities. Therefore, we investigated a simple subset V1 model of two-layer neurons coding the main features of interest in the study: orientation, contrast, spatial frequency (SF), and space. The model consists of orientation hyper-columns arranged into a hexagonal structure, with each hyper-column containing neurons responding to various contrasts and SFs. First layer neurons can be thought of simple cells whose responses are as follows:

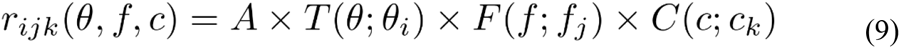

with “preferred” features (*θ_i_, f_j_, c_k_*) and the three normalized tuning functions to orientation, SF and contrast are described as wrapped-Gaussian (Swindale, 1998), log-Gaussian (Yuan et al., 2014) and hyperbolic ratio (Albrecht and Hamilton, 1982), respectively (*A* is the maximum amplitude of firing of the neuron). They are:

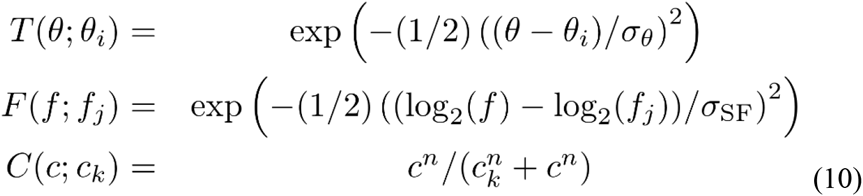

Remarque: for the contrast tuning, *c_k_* is the semi-saturation constant and can be called the “preferred” contrast of the neuron, since for contrasts around *c_k_* the neuron is the most informative above the input contrast(Chirimuuta and Tolhurst, 2005; May and Solomon, 2015a, b), and away from *c_k_* it asymptotes and provides no information about contrast input.

These simple cells feed a second layer of neurons through a spatial (excitatory center)–(inhibitory surround) connectivity structure, whose responses *R_ijk_* follow the conductance-based model (Grossberg, 1988; Piech et al., 2013)

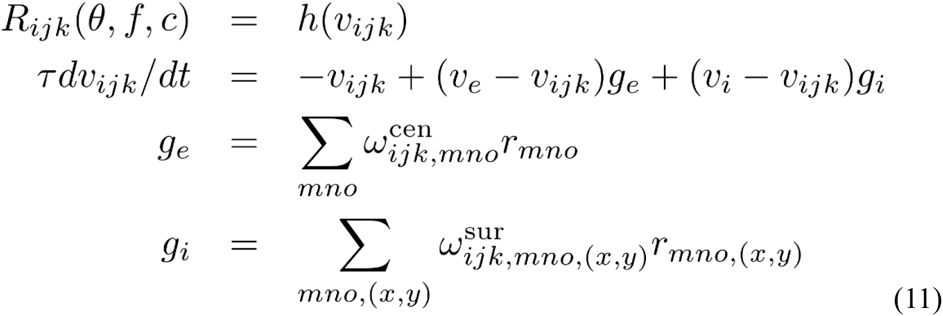

with *h()* a transducer (rectifying) function transforming voltage to firing rate and feature weights *ω*’s defined as:

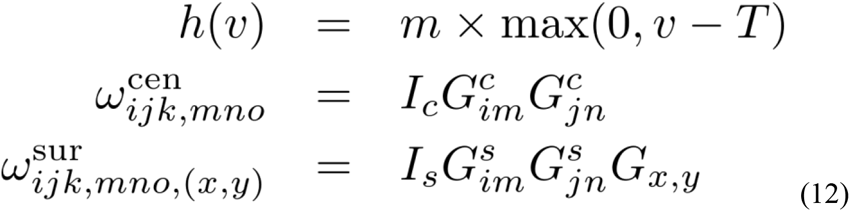

and the various parameters are: *T* is the voltage threshold of firing, *m* is the slope of voltage-to-firing rate relation, *τ* is the cell time constant, *v_e_* and *v_i_* are the excitatory and inhibitory equilibrium voltage potentials, *g_e_* and *g_i_* are respectively the excitatory and inhibitory conductances feeding the corresponding neuron through a weighted sum of first layer activities (*g_e_* sum is within hyper-column; *g_i_* sum is over all surrounding hyper-columns), *G_im_,j_n_* are Gaussian tuned feature weights (respectively within orientation and within SF; with possible different tuning widths indexed {*c, s*}), *G_x,y_* is a spatial weight function summing surrounding hyper-columns activity, and *I_c,s_* are the center/surround excitatory/inhibitory input strengths, respectively. Here, it is assumed that the weights are independent across features and iso-feature tuned (peaking at the receiving neuron preferred value (*i, j*), i.e. iso-orientation and iso-SF).

In the feed-forward model, equation (11) can be analytically solved, giving:

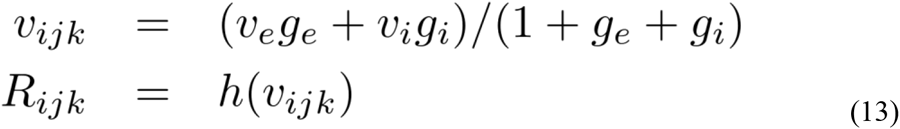

Using all relations above (Equations 9–13) and an input with uniform surround (all surrounding hyper-columns are stimulated with the same stimulus of orientation *θ_s_* and contrast *c_s_*), assuming the central stimulus (*θ_c_, c_c_*) excites the center hyper-column, the center input excitatory/inhibitory conductances can be analytically computed:

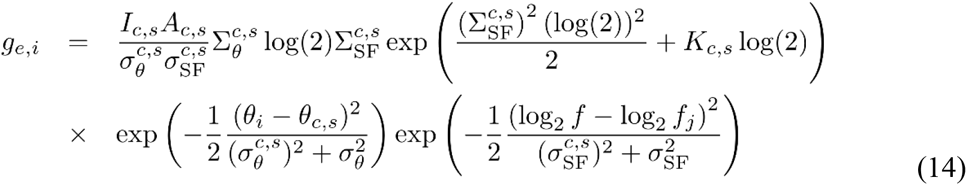

with the various constants defined as:

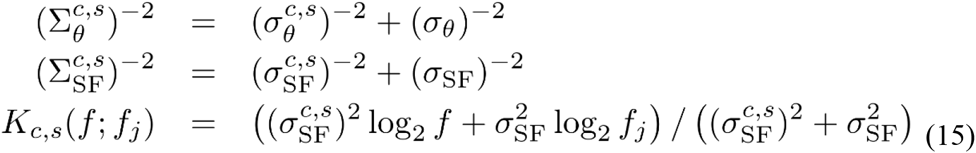

where 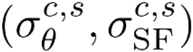 are the orientation and SF tuning widths of the weight functions *G_im_* and *G_jn_* (equation 12) for center-center and surround-to-center connections, respectively, *A_c,s_* are the contrast-weighted (*A ×C*(*c; c_k_*)) amplitude of firing of the input neurons *r_ijk_* for center/surround respectively, and *I_c,s_* are the excitatory and inhibitory inputs (cf. Equation (12)) with:

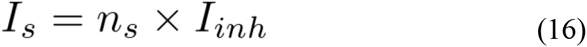

defining the total inhibitory input from all surrounding hyper-columns with mean inhibitory strength per hyper-column *I_inh_*, respectively, and *n_s_* is a “mean” number of surround hyper-columns influencing the center.

#### Fixed parameters and population model relations

In the model, a set of parameters were fixed based on previous literature. From Piech et al., we used a “normalized” conductance-based subtractive inhibition model with parameters: *g_thr_*=0.25, *g_xx_*=3/11, *g_yx_*=0.225, *v_e_*=14/3, *v_i_*=−2/3, *T_x_*=*T_y_*=1, *m_x_*=*m_y_*/2=1, *g_e0_*=*T_x_*/(*v_e_-T_x_*)-*g_thr_*, *g_i0_*=*T_y_*/(*v_e_-T_y_*), *v_0_*=*v_e_g_e0_*/(1+*g_e0_*) (*v_0_* is the activity for no input). Other parameters were: *n_s_*=6, *n*=2 (equation 6), *A*=2, *σ_SF_*=1, SF dimension sampling every ¼ octaves from ½ up to 64 c/d, orientation feature sampling every 2 degrees. The contrast tuning relation (equation (10)) was kept normalized by multiplying its amplitude by a factor (1+*c^k^_n_*).

#### Model based fitting of CSF and Tilt perception data

Here we investigate how based on the output activity *R_ijk_* of the network we can predict the perception of the subject in the two main features of interest, contrast detection for predicting the contrast sensitivity function and orientation identification for predicting the tilt repulsion effect. It is assumed that perception is based on decoding of the central hyper-column activities as described below.

Modeling the Contrast Sensitivity Function (CSF in 2D). In this experiment, the target stimulus is a vertical and uniform sine-wave grating limited in a circular spatial window, whose strength (contrast) is varied in order to measure the perception threshold across all SFs.

Given the uniform input stimulus, all input neurons *r_ijk_* of the central hyper-column plus the surrounding hyper-columns stimulated by the signal will have exactly the same input, and thus their activation across orientation, SF, and contrast will have the same profiles and peaks (for any 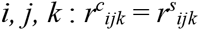). Second, given the task of detecting always a vertically oriented grating, by assuming that subjects disregard other orientated activities through an unspecified attentional mechanisms, we simplified into equation 10 the term over orientations into a value of one (in practice, this simplification can be thought of pooling these orientation neuronal activities into the constants *I_c,s_* or *A_c,s_*) and modeling only one orientation network activities.

In the above model description, one important function in predicting the contrast sensitivity at a given SF is the hyperbolic ratio of the neuronal population (Chirimuuta and Tolhurst, 2005; May and Solomon, 2015a, b), which for contrast detection and a given SF can be assumed to be the neurons with the smallest semi-saturation constant 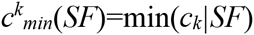. In our model, to predict the CSF across SFs, we additionally need to properly describe the relation between 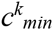 and SF together with SF tuning width versus preferred SF. Based on previous neurophysiological reports about neuronal sensitivity function (Anzai et al., 1995; Kiorpes et al., 1998), we fixed the neuronal population minimum sensitivity to follow the inverse CSF shape of equation:

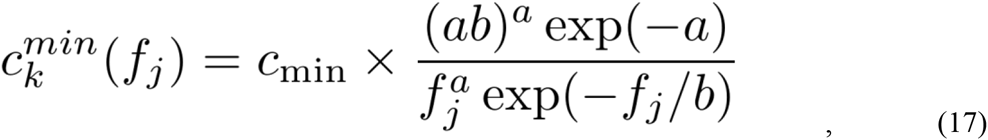

and for the SF tuning width (Albrecht and Hamilton, 1982) as a function of preferred SF, the following relation:

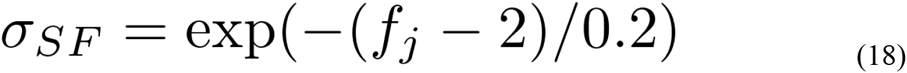

where *f_j_* is the preferred SF of the neuron and parameters (*a, b, c_min_*) define the neuronal sensitivity function (NSF) across SFs. We fixed 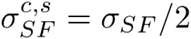 because of center/surround tuning widths entanglement in predicting behavioral results, and *I_c_*=1 and *n_s_*=6, and neurons with *c_k_*>1000 were pruned.

Last step, for predicting the contrast sensitivity function across SFs, we used the standard signal detection theory providing us the psychometric function:

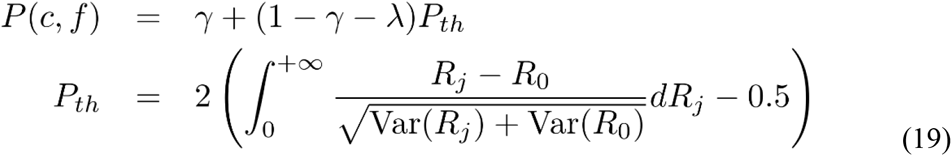

with the lapse and guess rates obtained as described in the above section ***Data Analysis***, and *R_0_* being the activity of the neurons for no signal input.

From the above model description, there are only four free model parameters that need to be adjusted for predicting the full CSF: *c_min_, I_inh_, a, b*. It was done by replacing equation (5) with equation (19) and following all remaining steps.

#### Modeling Orientation Identification (Tilt illusion)

For this feature, we make a different set of simplifications in the model feature space based on the experimental design for tilt perception. Here, center and surround hyper-columns are stimulated with varying orientations while the contrast of the center and surround stimuli are kept constant and equal. Therefore, we can describe the two-layer neuronal network activities through the above mathematical development (equations (9)–(16)). But here, we fixed the input layer contrast activation at *A*=2 and *c*=1 (near maximum contrast(Piech et al., 2013)), and given the task of identifying the orientation of the center stimulus for a fixed SF, we simplified into equation (14) the term over SFs by assuming subjects disregard other SF neuronal activities(Georgeson, 1973) (e.g. through an unspecified attentional mechanism), giving:

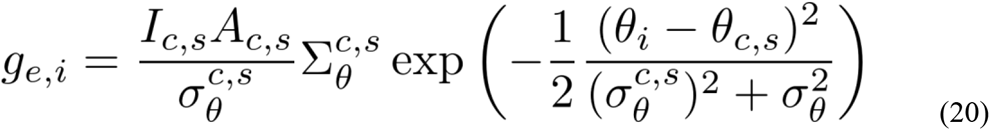

with all constants defined in model description section. We fixed the center and surround feed-forward orientation tuning widths to 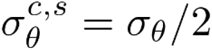.

Last, to predict the orientation psychometric function (tilt data), we decoded the perceived orientation (*a* in equation (4)) of the stimulus as the orientation vector average of the central hyper-column activities(Gilbert and Wiesel, 1990a; Seung and Sompolinsky, 1993; Tzvetanov, 2012), while the discrimination threshold was left as independent parameter(Fu et al., 2010).

### Neurophysiological data re-analysis

The direction of motion tuning curves were fit with a wrapped von-Mises function of the form:

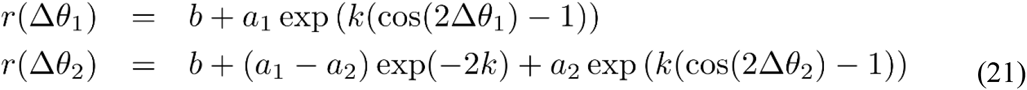

where r is the response of the neuron, and the parameters *b, a1, a2*, and *k* allow to compute the physiological parameters of interest, minimum firing rate, the two amplitudes to opposite direction of motions, and half-width at half-amplitude, as:

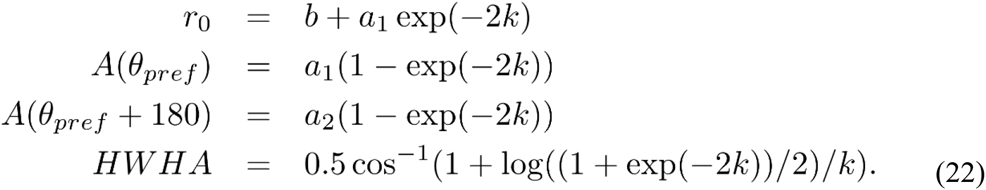

The two variables *Δθ* are defined as *Δθ1=θ-θ_pref_* and *Δθ2=θ-θ_pref_−180*, each defined in the range ±90 degrees, and the additional term for *r* (*Δθ_2_*) allows continuity at the boundary. The fitting was done by minimizing the *χ*^2^ between the data and the model function, and because standard errors were not present, it was assumed *sei* = (FANO×*y*_i_/5)^½^ with FANO=1.5, and for the few cases whereyi=0, sei=0.5477. We performed three successive fits: (1) an orientation tuned function fixing *a*_1_=*a*_2_ (4 free parameters: *b, a, k, θ_pref_*); (2) a direction tuned function leaving parameters *a*_1_ and *a*_2_ independent; and (3) a flat-topped version of the equation^54^ allowing for more peaked or flatter types of tuning curves, that included one more parameter *v* (in equation 21, *Δθ_1,2_* is replaced by *Δθ_1,2_*/2+ν sin(*Δθ_1,2_*)). Multiple refits were done with random starting points for finding the best set of parameters. Each fitted curve was then used to test, with a nested F-test (Tzvetanov, 2016), whether it described the data better than the global mean of the data, i.e. whether the cell could be described as tuned. Then, if two or more functions described the data better than the mean, we further used a nested F-test to test whether the more complex functions (with additional parameters) described the data better than the simpler one, and selected the corresponding model. In the models with different amplitudes to opposite motion direction, the amplitude of firing was defined as the higher of the two. In the flat-top model, the *HWHA* was directly searched in a discretized direction space (0.1 degrees steps). At the end, we additionally eliminated any fitted curve with too narrow of tuning (HWHA<15), which discarded 8 old cells and 7 young cells. Two example fits are presented on Figure 7a, b for one orientation and one direction tuned cells, together with their orientation and direction bias values.

### Neurophysiological Experimental Materials and Methods

Materials and methods are same to Yu Fu, et al., 2013 Cerebral Cortex(Fu et al., 2013). Briefly:

#### Animals

Subjects for this study were 14 male rhesus monkeys (Macaca mulatta) classified into 2 groups: the young adults group consisted of 4 monkeys who were 5.80 ± 0.50 (mean ± standard deviation) years old; the old group included 10 senescent monkeys at ages of 25.46 ± 1.90 years. Monkeys were examined ophthalmoscopically to exclude possible optical or retinal problems that would impair visual function.

#### Animal Preparation and Recording

Subjects were sedated with Ketamine HCl (10–15 mg/kg, i.m., ParkeDavis, Morris Plains, NJ, and USA) and then anesthetized with 3–5% halothane (Halocarton Laboratories, River Edge, NJ, USA) in a 70:30 mixture of N_2_O: O_2_. Intravenous and tracheal cannulae were inserted. Are V1 was exposed by a craniotomy surgery. The anesthesia and paralysis were properly maintained during whole experiment processing. Multiple vital life signals including retina condition were carefully monitored and kept in a constant level.

#### Visual Stimulation

For each isolated single unit, the dominant eye and cell’s receptive field were anchored before each neural recording. To quantify the orientation and direction selectivities of V1 cell, drifting bars were used, whose width, length, and moving speed were adjusted to elicit strongest response from the recorded cell. The direction of motion of each presented bar was orthogonal to its orientation. We used moving bars at 24 randomly chosen movement directions, ranging from 0° to 360° in steps of 15° to compile the tuning curves for the cells studied. Each direction was presented 10 times. The inter-trial interval was 2–5s. The luminance of the stimuli used was 39 cd/m^2^ for white and 0.95 cd/m^2^ for black.

#### Data Collection

Data Collection and Analysis Signals recorded from the microelectrode were amplified (1000×), band-pass filtered (300–10000 Hz), and then digitized (sampling frequency of 20 kHz) by using an acquisition board (National Instruments, Austin, TX, USA) controlled by IGOR software (WaveMetrics, Portland, OR, USA). Such original signals were stored in a computer for offline analyses. The responses to moving bars were defined as the maximal value in the peristimulus time histogram (PSTH, bin width of 10 ms) during the stimulation period. Before each drifting bar was presented, the spontaneous (baseline) activity was recorded during a period of 0.5–0.7 s while the screen with average luminance was presented.

## Results

### Contrast sensitivity functions

Contrast sensitivity function (CSF), representing the inverse of the minimum detectable contrast at each spatial frequency (SF), was measured for each subject prior to the tilt illusion measurement (Figure 2a, c and d). The contrast sensitivity (CS) was significantly modulated by SF (F (10, 380) = 599.56, P= 0.0001) and CS of the elder group was significantly lower than the young adults group across all spatial frequencies (F (1, 38) = 7.44, P=0.01), confirming previous reports(Owsley, 2011). The interaction of the SF and groups was also significant (F (10, 380) = 7.23, P= 0.01). From each individual CSF, we chose two spatial frequencies (SFs), one near the peak sensitivity (low-SF) and one higher (high-SF) (Figure 2c, squares), with the condition that the sensitivity at the high-SF was not too low for allowing stimulus perception at next stage of measures of center-surround tilt illusion.

### Tilt illusion measurement

Perceived verticality of the target grating under center-surround orientation differences of 0, ±15, ±30 and ±75 degrees for each subject was measured at low- and high-SF (Figure 2b). The amount of orientation misperception for both groups was systematically modulated by surround orientations (SOs) and SFs (Surr. Or.: F (3,114) =240.41, P=0.0001; SF: F (1, 38) =75.14, P=0.0001) (Figure 2e), consistent with previous reports (Georgeson, 1973; Wenderoth and Johnstone, 1988; Smith and Wenderoth, 1999). Importantly, analysis limited to ±15° and ±30° SOs, corresponding to the direct form of the tilt illusion (Wenderoth and Johnstone, 1988; Smith and Wenderoth, 1999; Wenderoth and Smith, 1999), repulsion, revealed that perception through the elder visual system exhibited much stronger tilt repulsion effects across all measured SFs in comparison to the younger adults (F(1,38)=6.62, P=0.014) (Figure 2e). The amount of misperception at ±15° SO was significantly different than that at ±30° (F (1, 38) =53.79, P=0.0001). There was no significant interactions between SOs and groups (F (1, 38) =0.32, P=0.57) indicating similar young-elder differences at both SOs. The amount of misperception for low- and high-SFs across groups was significantly modulated (F (1, 38) =89.72, P=0.0001), but there was no interaction between SFs and groups (F (1, 38) =1.56, P=0.22) that suggested same effects of SFs across age groups. There was a significant interaction between SOs and SFs (F (1, 38) =41.28, P=0.0001), while there was no significant interactions among SOs, SFs and Groups (F (1, 38) =0.15, P=0.70), again showing no differences of effects across age.

Orientation discrimination thresholds in elder group were larger than younger adults (F (1, 38) =7.25, P=0.01), and they were modulated by SOs (F (3, 114) =11.86, P=0.0001) and SFs (F (1, 38) =49.66, P=0.0001). There was also a main interaction between SOs and SFs (F (3, 114) =7.58, P=0.001) showing a different trend of threshold variation with surround orientation at different SFs (Figure 2f). All other interactions that included age group were not significant (SOs × Groups: F (3,114) =1.73, P=0.17; SFs × Groups: F (1, 38) =1.04, P=0.32; SOs × SFs × Groups: F (3,114) =1.73, P=0.17).

It is possible that the different high-SF used in younger and elder groups contributed to the differences in misperception between them. We found, as in previous work (Smith and Wenderoth, 1999), that the amount of orientation bias was significantly related to the SF under both ±15° and ±30° SOs conditions for both elder and younger groups (Spearman rank’s correlation; the elder: r=0.58, P=8.5*10^−5^ at ±15°, r=0.39, P=0.013 at ±30°; the younger adults: r=0.74, P=3.6*10^−8^ at ±15°, r=0.63, P=1.2*10^−5^ at ±30°) (Figure 3a, b). This suggested that adjusting for the variation of tilt illusion with SF (Smith and Wenderoth, 1999) still showed the elder exhibited stronger misperception. Additionally, the elder group had globally lower high-SF (t (38) =5.37, P=0.0001, unpaired t-test) than the younger group, showing that their stronger repulsion effects cannot be explained by this experimental manipulation.

**Figure 3:**
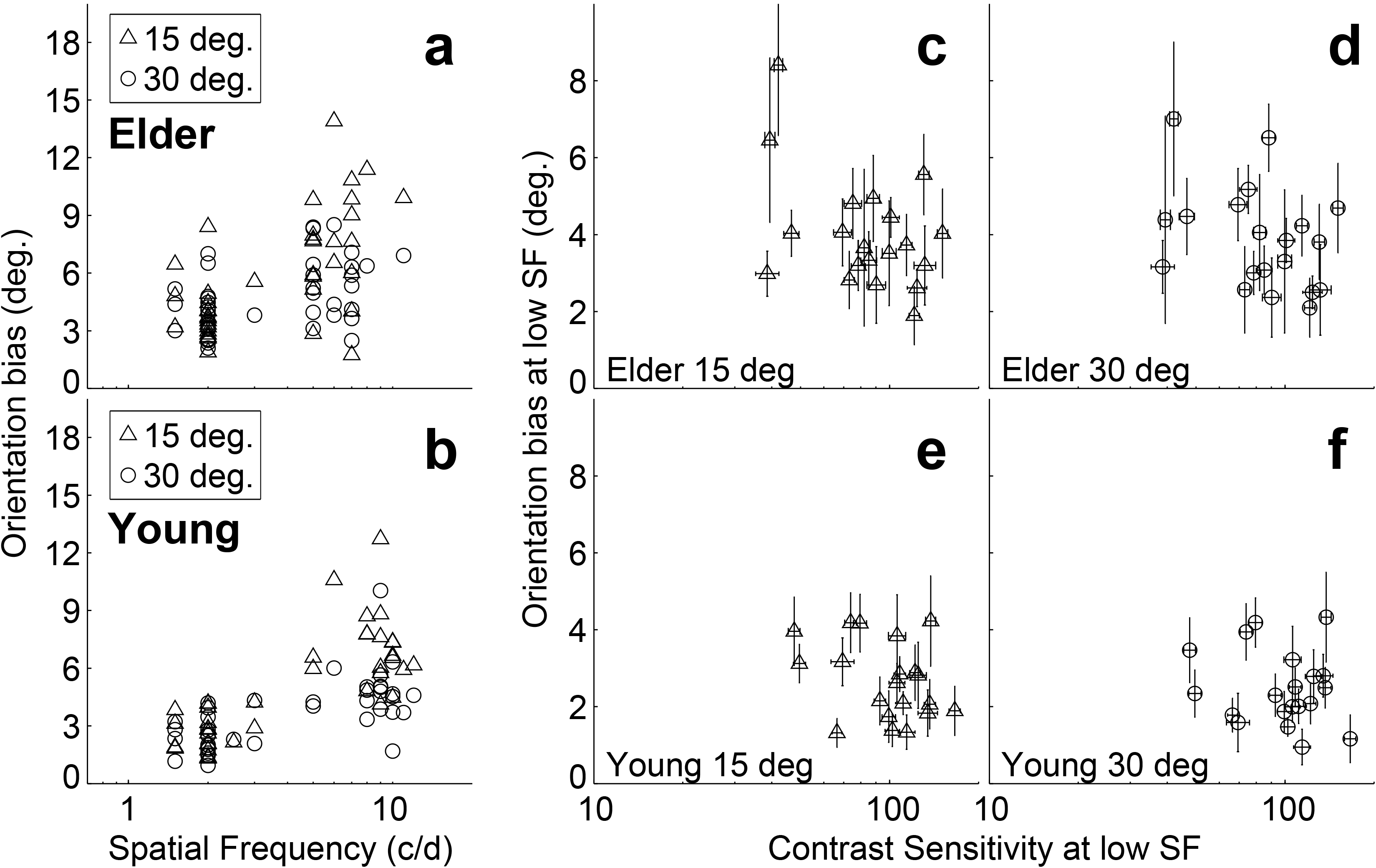
Relation between tilt repulsion, SF and CS at low-SF. Relations between tilt repulsion and SFs under ±15° (triangles) and ±30° (circles) SOs across all measured subjects in the elder (a) and younger (b) groups; Relations between tilt repulsion and CS at low-SF in the elder group (±15° in (c); ±30° in (d)) and younger (±15° in (e); ±30° in (f)) groups. Error bars are bootstrapped SE.

### Relation between tilt illusion bias and contrast sensitivity

The elder group exhibited decreased contrast sensitivity and increased amount of tilt illusion compared to the younger adults. To reveal any possible relation between tilt repulsion and peak CS, we correlated tilt and CS measured at low-SF condition, where the SF values for both groups are similar. The results showed that there was no significant relation between bias and CS within each age group (the elder: r=−0.25, P=0.29 at ±15°, r=−0.36, P=0.12 at ±30°; the younger adults: r=−0.21, P=0.38 at ±15°, r=−0.02, P=0.92 at ±30°) (Figure 3c-f).

### Relation between tilt illusion bias and contrast suppression ratios

A recent report demonstrated stronger surround suppression in the elder when compared to young population (Karas and McKendrick, 2015). Using our stimuli, a center-surround contrast discrimination task was conducted for all participants in order to investigate how it correlated to our own findings with the tilt repulsion. In our results, the suppression ratios were lower than 1 for both groups (Fig. 4), and the elder’s ratios were significantly lower than the younger adults (t (38) =4.48, p<0.0001), thus replicating their finding (Fig. 7 in Karas & McKendrick, 2015). If the two psychophysical measures probe the same underlying visual system, we should expect a correlation between them. To analyze this relationship, the expected orientation bias at 4 c/d (Bias (SF=4)) for each individual subject was extracted from a line regression between orientation bias and log-SFs (Fig.3a, b). There was a significant negative correlation in the younger group at both surround orientations (Figure 4a: r=−0.77, p<0.0001 at 15°; r=−0.64, p<0.001 at 30°) and in the elder group at 15° (Figure 4b: r=−0.51, p<0.05 at 15°; r=−0.25, p=0.29 at 30°).

**Figure 4:**
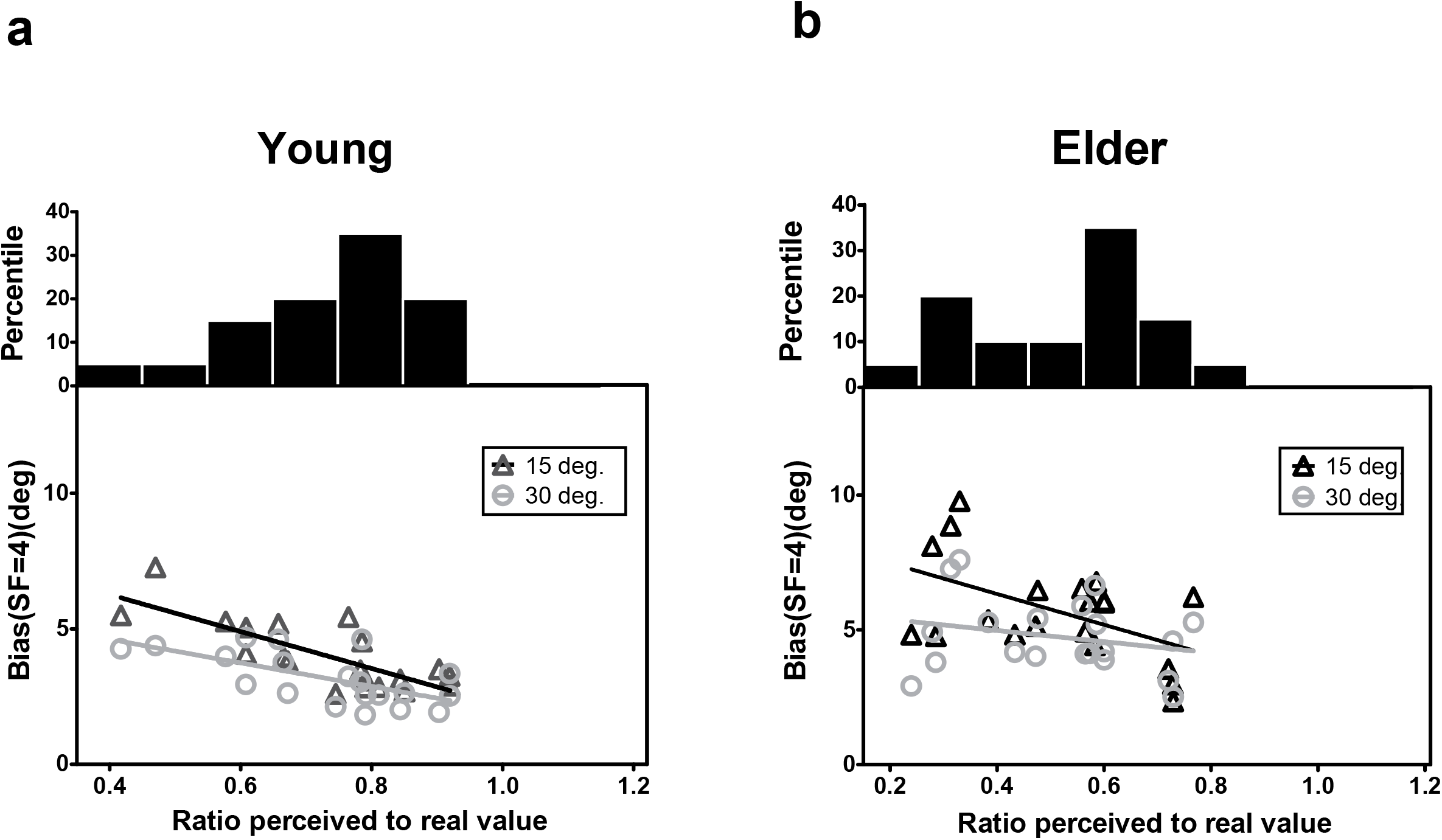
Relation between tilt repulsion and contrast suppression ratio. Correlation between tilt repulsion (bias, at spatial frequency equals 4 c/d) and contrast surround suppression ratio (ratio of perceived contrast/physical reference contrast) for younger (a) and elder adults (b).

## Modeling

To further our understanding of early visual processing and the plausible underlying network changes with aging, based on the learned knowledge that neuronal activity in V1 could be the substrate of orientation identification and detection(Georgeson, 1973; Gilbert and Wiesel, 1990a; Kapadia et al., 2000; Tzvetanov, 2012), we modeled tilt repulsion effects and contrast sensitivity including the spatial lateral inhibition in V1.

While very appealing, psychophysical modeling based on the V1 neuronal responses relates each behavioral variable to different tuning characteristics: orientation misperception is obtained through orientation tuning and center-surround inhibitory interactions (Figure 5a-c), while CSF could be obtained through the SF and contrast tuning of the neurons (Figure 5d-g).

**Figure 5:**
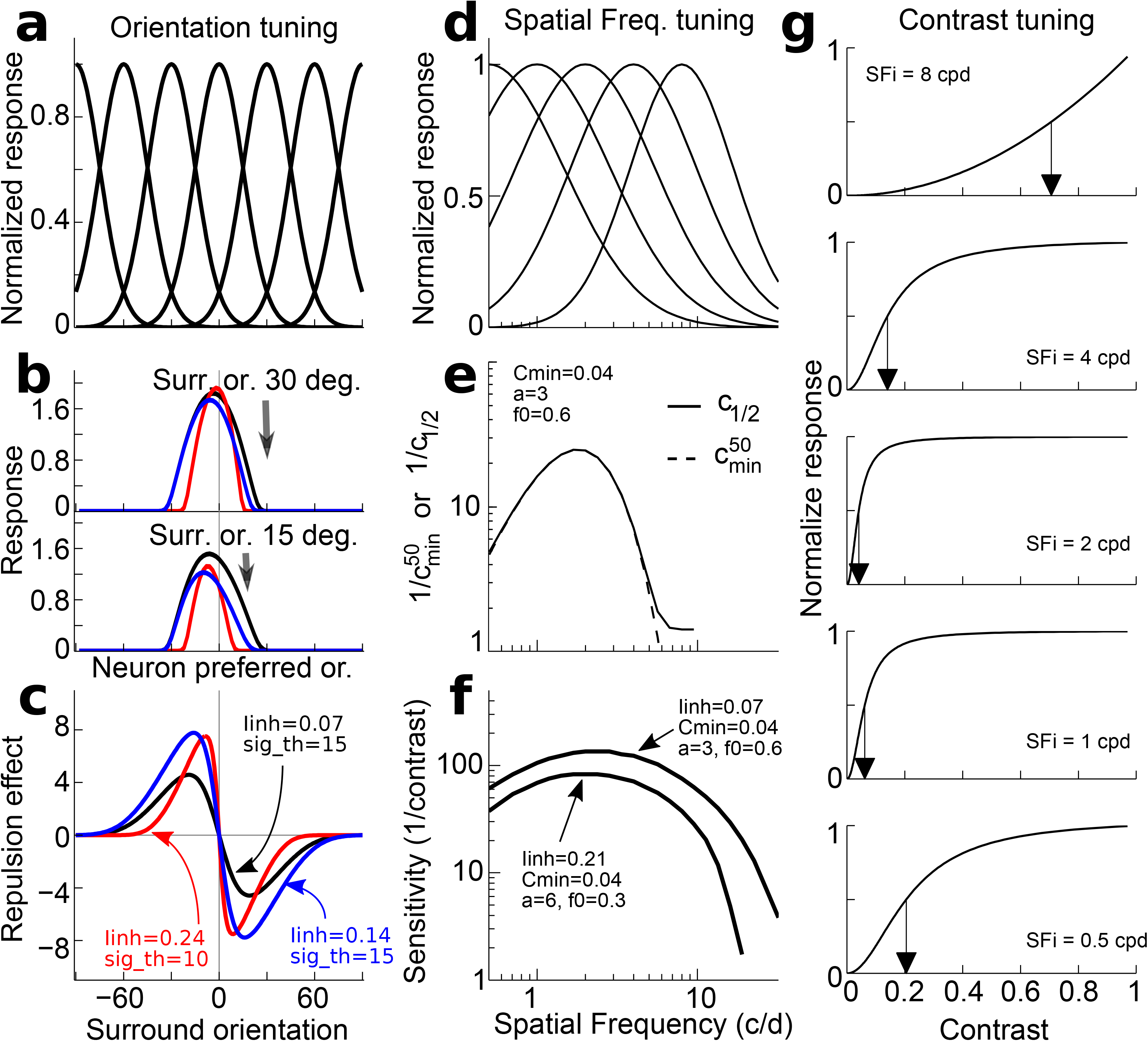
V1 model illustration. (a-c) Example and prediction for orientation coding and decoding. (a) Uniform orientation tuning of the neuronal population. (b) Response of the neuronal population to center of 0 deg. and two different surround orientations of ±30° (top) and ±15° (bottom). (c) Orientation prediction of the model from the population responses for various surround orientations. Examples for three different set of parameters in red, black and blue. (d-g) Example and prediction for (SF) and contrast tuning coding and decoding. (d) SF tuning examples, with the characteristic tuning width decrease with increasing preferred SF. (e) Example of the relation between the minimum contrast semi-saturation constant and preferred SF. (f) Examples of CSF prediction for two sets of model parameters. (g) Examples of contrast response functions in the model for the minimum semi-saturation constant at few preferred SFs (arrows depict half-amplitude constant).

To obtain an insight into the differences between the elder and younger adults (for instance, lateral inhibition, orientation tuning, contrast tuning or a combination of them), we modeled a two-layer feed-forward surround-to-center inhibitory network of V1 neurons. The first layer cells had generic tuning characteristics coding orientation (*θ*), spatial frequency (*f*) and contrast (*c*) (e.g. simple cells; Figure 5a, d, g). Their responses *r_ijk_* can be described as:

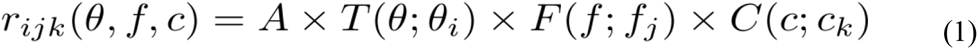

with *T()*, *F()* and *C()* being the neural tuning functions to orientation, spatial frequency and contrast, respectively. This first layer neurones feed a second layer of cells through a spatial excitatory-centre and inhibitory-surround connectivity structure, whose responses *R_ijk_* follow a conductance-based model (Grossberg, 1988; Piech et al., 2013):

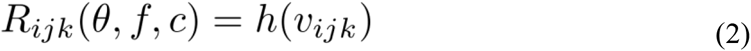

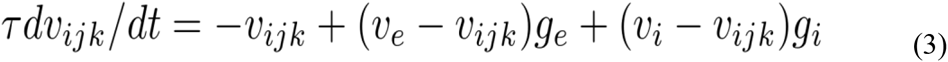

where *h()* is a transducer function transforming voltage to firing rate, *τ* is the cell time constant, *v_e_* and *v_i_* are the excitatory and inhibitory equilibrium voltage potentials, and *g_e_* and *g_i_* are respectively the total excitatory and inhibitory conductances feeding the corresponding neuron through a weighted center-surround sum of first layer activities. In this model, the tilt illusion and contrast sensitivity function (Figure 5c, f) are related through a single parameter, the amount of surround-to-center inhibition (defined as *I_inh_*). The amplitude and shape of the orientation misperception is dependent on *I_inh_* and orientation tuning width (defined as *σθ*) (Figure 5a-c). The CSF is dependent on *I_inh_* (because the grating stimulus excites center and surround mechanisms) together with the smallest contrast semi-saturation constant (defined as *c_min_*) and its relation to the SF tuning (Figure 5d-g).

We fitted the model first to tilt perception data (two free parameters: *I_inh_, σθ*), and then to CSF data by using lateral inhibition as a fixed parameter (for CSF prediction, there were three free parameters: *c_min_, a, b*; with *c_min_* corresponding to the best neuronal contrast sensitivity across all SFs; e.g. Figure 5e, peak value; see Methods for fitting details). The model fit to the tilt repulsion data of ±15 and ±30 degrees provided independent estimates of orientation tuning widths and surround to center inhibitory amplitudes at both low- and high-SFs for each subject (Figure 6a, b). Age differences in tilt repulsion near the peak of perceptual sensitivity (low-SF) were explained by stronger lateral inhibition in the aged group (t (38) =2.95, P=0.0054, log-transformed values), while no differences in orientation tuning widths were observed (t (38) =1.25, P=0.22). Additionally, both parameters co-varied with the SF at which it was measured across all population: lateral inhibition increased with higher SFs (r=0.65, P=4.4*10^−11^, n=80), while neuronal tuning widths tended to decrease with higher SFs (r=−0.31, P=0.0053, n=80). This last finding resonates with the established physiological findings of sharper orientation tuning widths for cells with higher preferred SFs (Tolhurst and Thompson, 1981; De Valois et al., 1982), and provides strong support for our modeling approach. The model fit to the CSF data provided the minimum semi-saturation constant of the contrast response function near the peak of the CSF. Age group did not show differences in this optimal neuronal contrast sensitivity (t (38) =−0.19, P=0.85, log-transformed values (Figure 6c), but instead strongly correlated with the peak contrast sensitivity of the subjects (young group: r=−0.67, P=0.001; old group: r=−0.51, P=0.022). On the other hand, lateral inhibition was not correlated with peak contrast sensitivity (young: r=−0.27, P=0.24; old: r=−0.37, P=0.11) (Figure 6d).

**Figure 6:**
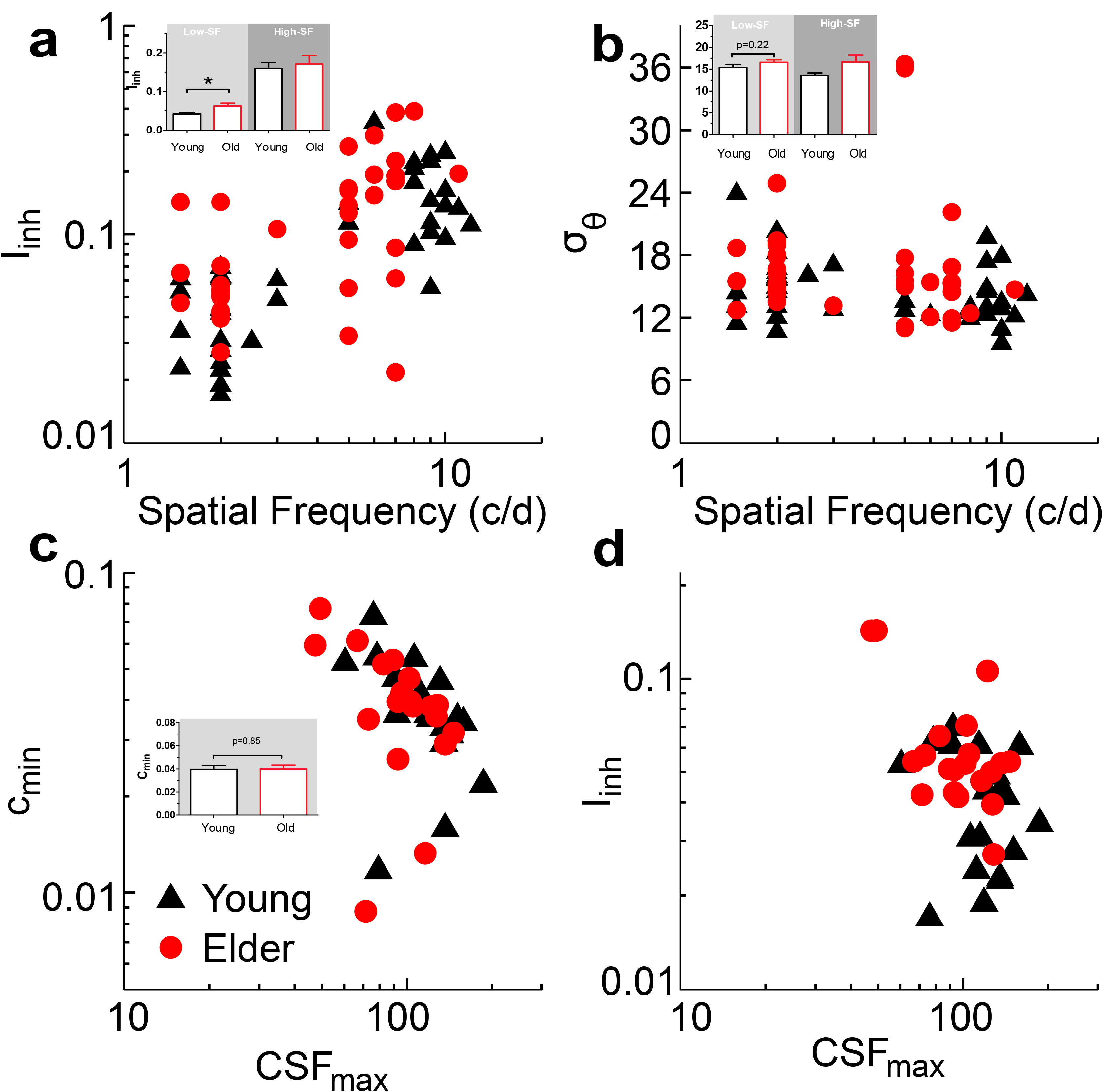
Model fitting results. Summary plots of subjects model parameters (Iinh, σ_θ_, c_min_) obtained from tilt repulsion data (a, b) and contrast sensitivity data (c) and relations to the SF (a, b) or peak contrast sensitivity (c, d). Insets depict histograms of the corresponding variable on the ordinate with significance values for the low-SF case.

This combined modeling of orientation identification and contrast detection shows that neuronal contrast sensitivity and lateral interactions are not straightforwardly associated to the perceptual outcome and need simultaneously to be considered. Aging affected the strength of lateral interactions while individual subjects differences in neuronal contrast sensitivities still reliably represented their individual perceptual sensitivities.

### Neurophysiological data re-analysis

From the above modeling results, we found that orientation tuning widths are not systematically changed with aging, which is consistent with the psychophysical reports proposing that orientation discrimination is maintained in older population (Delahunt et al., 2008; Govenlock et al., 2009). But these behavioral results are not directly comparable with our published neurophysiological evidences from monkeys and cats, showing that orientation processing in the elder primary visual cortex is substantially affected (Leventhal et al., 2003; Hua et al., 2006; Fu et al., 2010; Fu et al., 2013).

These previous analyses did not consider the orientation tuning width of neurons. Instead, it was obtained through the method of measuring orientation or direction tuning of the cells and analyzing them through the orientation and direction bias indexes (OB/DB). It is important to note that these indexes are difficult to interpret with respect to changes in tuning properties (background, amplitude, or tuning width, see (Mazurek et al., 2014; Tzvetanov, 2016)), because of two important points. Firstly, the analysis based on them includes all neurons sampled during the study, even the ones that have a circularly flat response (i.e. they respond uniformly to all directions of the stimulus). Thus, a decrease/increase in OB/DB could reflect the fact that there are less/more tuned cells in the first condition when compared to the second, simply because the OB/DB of these not tuned cells is close to zero (but not zero). Secondly, even for the cells that have clear orientation/direction tuning, the decrease/increase of OB/DB does not provide information about what changed in the tuning property of the neurons. As a simple example, consider a theoretical Gaussian orientation tuned cell with minimum firing rate (r0) of 10 Hz, amplitude (A) of 40 Hz, and σ of 25 degrees (giving HWHA of about 29.4 degrees). Its OB value, when sampled every 1 degree, is 0.40. If a second cell has simply a higher minimum firing rate of 30 Hz, and all other parameters and sampling are identical, it provides a value of 0.22 for its associated OB (but the other parameters also influence the final OB/DB estimate, see(Mazurek et al., 2014; Tzvetanov, 2016)). This demonstrates that a change of OB/DB alone is not informative about the nature of the underlying modification, whether there are less cells coding for orientation/direction or the parameters of the tuned cells changed.

Re-analysis of these data through tuning curves allowed to separate effects onto the parameters of tuning: tuning width, minimum firing rate, and amplitude, which provided a clearer picture about age-related changes in orientation tuning properties.

Here, we re-analyzed one data set (Fu et al., 2013), which measured the tuning curves with bar stimuli, containing 264 cells from 10 senescent monkeys and 140 cells from 4 young monkeys, by fitting direction/orientation tuning functions to each neuronal data set (Figure 7a, b). Using these fitting results, we first tested, with an F-test, whether a given neuron’s data can be considered as tuned to orientation/direction of motion (Mazurek et al., 2014; Tzvetanov, 2016). Then, from the tuned cells, we extracted three important parameters of the tuning curve: the minimum firing rate in the tuning curve (*r*_0_), amplitude of firing (*A*), and tuning width as half-width at half-amplitude (HWHA). First, in the elder monkeys, 143/264 (54.17%) neurons were found as tuned, a proportion which significantly differed (*χ*^2^ = 5.014, P < 0.05) from the 92/140 (65.71%) tuned cells found in the young animals. This confirmed the previous reports of smaller number of cells with orientation/direction coding capacities in the primary visual cortex of senescent animals (Schmolesky et al., 2000; Leventhal et al., 2003; Hua et al., 2006; Fu et al., 2013). Second, *r_0_*, (young: 32.83±1.80; old: 52.11±2.68), which here we consider as neural noise, was substantially increased (t(226.51)=5.28, P<0.001) while amplitude *A* (young: 79.47±4.60; old: 77.66±4.29) and tuning width *HWHA* (young: 35.73±1.37; old: 33.85±1.00) were similar (A: t(233)=−0.28, P=0.78; *HWHA:* t(233)=−1.13, P=0.26) in the two age groups of neurons (Figure 7c-e). Additionally, the maximum firing rate of the cells (*r_0_+A*) in the two populations (young: 112.30±4.86; old: 129.77±4.88) differed (t (233) =2.42, P<0.05) (Figure 7f), which was mainly attributed to the change in *r_0_*.

**Figure 7:**
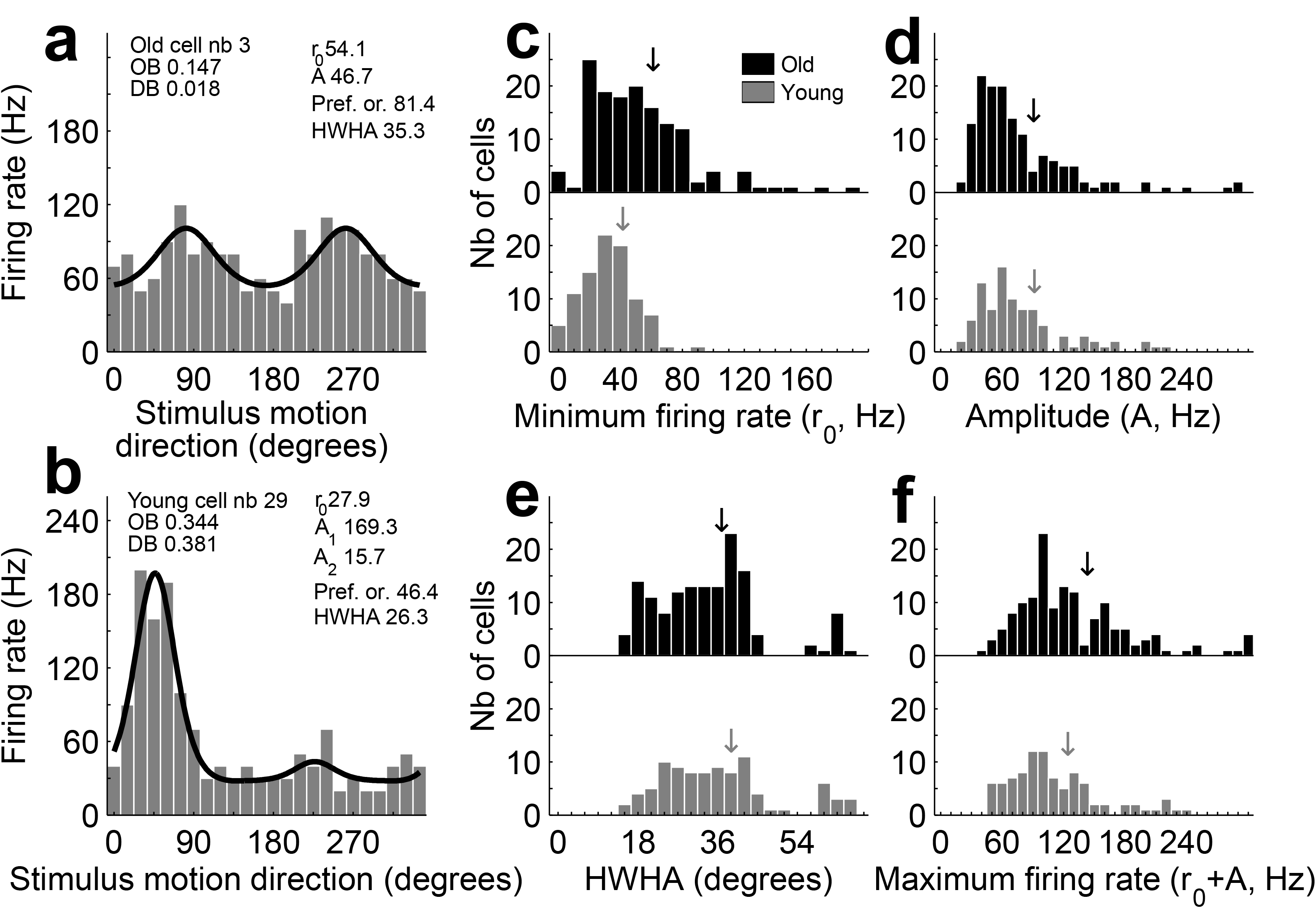
Neurophysiological data re-analysis of orientation tuning. Examples of fitting results for orientation (a) and direction (b) of motion selective cells; OB – orientation bias index, DB – direction bias index. Distribution of minimum firing rate (c), amplitude (d), *HWHA* (e) and maximum firing rate (f) for the young and old cells. Arrows depict mean values.

Since this data set was not analyzed for possible presence of multi-unit activity (MUA), we considered that the tuning curves might be contaminated by multiple closely tuned cells. While we would expect this effect to be similar between the old and young cells because of the same methodology employed within the study and across monkeys, we decided to check for plausible contamination by restricting the main analysis to a subset of the cells with the best tuning widths. The idea is that these cells should be much less contaminated since they have the narrowest tuning widths. For that purpose, we selected the 30% cells with smallest tuning widths in each population (for old, n=43; for young, n=28) and re-applied the main text comparisons on the three tuning properties. The minimum firing rate r0 (young: 36.0±3.9; old: 54.1±5.7) was significantly different between the two populations (t (67.59) =2.61, p=0.011), while the amplitude A (young: 100.9±9.9; old: 89.7±9.6) and HWHA (young: 22.3±0.7; old: 20.9±0.6) were both not significantly different between the two cell types (A: t (65.0) =−0.81, p=0.42; HWHA: t (59.7) =−1.61, p=0.11). This restricted sub-sample analysis confirms the main text results, but on those cells that should not be considered as strongly affected by MUA.

Here, we also considered whether the fit quality could be different between the two categories of cells, that is, we checked whether the tuned old cells might have worse tuning properties than the young cells through a goodness-of-fit measure. For that purpose, we first computed the classical coefficient of determination, r^2^, in both populations. The old cells had globally higher r^2^ (Wilcoxon rank sum test for differences in medians: p=0.026). The above found increase in minimum firing rate (r0) of old cells provides globally higher firing rates for the old cells thus artificially inflating the total sum of squares and creating the impression of better model description of that data set. Therefore we sought a more appropriate way by computing for each cell the likelihood to obtain the given data set from the best fitted function based on the Poisson statistics for each datum, and re-transformed it into individual cell p-value by taking the likelihood to the power of (1/24). Once the variability is properly accounted for, the young and old cells did not show significant differences in goodness-of-fit (Wilcoxon test: p=0.31).

The key results described above were also confirmed, out of the amplitude effect, by re-analyzing a second data set (Fu et al., 2010) from our laboratory about orientation/direction tuning and surround effects onto CRF. In their article, Fu et al. analyzed possible changes with aging of surround suppression onto the orientation tuning properties of the classical receptive field (CRF)(Gilbert and Wiesel, 1990b) measured in young and old monkeys (4 young monkeys mean age 5.5 years and 3 old monkeys with mean age 28 years; for further methodological details please refer to the article). They performed it by presenting luminance sine gratings into the CRF first for characterizing the standard orientation/direction tuning, and then in one second condition, they presented the optimal center orientation in the surround and re-measured the orientation tuning of the cell. The results were interpreted and analyzed with respect to the old-fashioned orientation bias/direction bias (OB/DB) indexes, which again introduced unwanted interpretation pitfalls (Mazurek et al., 2014; Tzvetanov, 2016).

Here, we first re-analyzed the main CRF orientation/direction tuning parameters. In the old cells, 20 out of 46 cells were found tuned while in the young cell population 66 out of 81 were tuned; this difference was highly significant (Z=4.4, p<0.0001). The minimum firing rate r0 in the old cell population (61.3±4.7) was significantly higher than the young cells (30.6±2.6)(Wilcoxon rank sum test: Z=4.8, p<0.00001); the amplitudes A in the old cells (37.1±4.1) were found significantly lower than those of the young cells (63.7±4.4)(Z=−3.08, p=0.0021); HWHA were not different between old (44.5±3.2) and young cells (41.2±1.6)(Z=1.21, p=0.23); and the total amplitude of firing (r0+A) was similar between old (98.4±6.5) and young (94.3±0.6) cells (Z=1.10, p=0.27). Overall, this data set confirms the findings of minimum firing rate change and no tuning width change, while the amplitudes seem to be differentially modulated by the type of stimulus used for the measures.

This data set was also interesting because the authors directly investigated in old and young cells the tuning with the presence of optimal surround orientation, known to inhibit the activity of the center. We also fitted this condition with the orientation/direction tuning functions. In this condition, we discovered a puzzling behavior of the cells orientation/direction tuning, especially in the old cells. From the 46 old cells only 14 cells were found tuned in this condition, and from those, only 6 cells were simultaneously tuned in the standard CRF tuning measure (previous paragraph). That is, we found that 8 cells “popped-out” as being tuned with the presence of the surround while they were not in the standard measure (no surround). For the young cells, 35/81 were orientation tuned in the surround condition, and among them only 2 cells were not tuned in the standard condition. These results are partly consistent with the original article conclusion based on the OB/DB indexes (orientation bias/direction bias) that there are even smaller number of orientation/direction tuned cells when surround is present, but also with the main finding in the previous paragraph about the tuning parameters r0 and HWHA (means±s.e. for surround condition; r0: young – 36.2±3.1, old – 64.8±6.4; HWHA: young – 37.1±2.1, old – 42.0±4.1), but not for the amplitudes A (young: 42.7±3.9; old: 35.8±4.8).

We would like to emphasize that the apparent inconsistency of the original interpretation of “decreased strength of surround suppression” in the cells of senescent animals with the re-analysis presented here is due to the old-fashioned way of analysis through the OB/DB indexes or other composite variables (Mazurek et al., 2014; Tzvetanov, 2016), which are obtained from a combination of tuning parameters.

On the contrary, one has to use a clear statistical test for presence of tuning for each individual cell and then analyze the tuned cells parameters by discarding from the analysis the cells that are not tuned. In the current analysis, from the statistical test and tuning parameters changes, the exact nature of stronger/weaker suppression in the older cells seems to us difficult to ascertain for multiple reasons. First, in both old and young cells, the number of tuned cells when the surround was present was decreased, leading to the possibility of stronger surround suppression in both populations, which destroys orientation/direction tuning of the cell. Second, the mean amplitude of firing of the tuned cells with surround present seemed, when compared to the CRF tuning, unchanged in the old cells but lower in the young cells, which leads to the possibility that old cells might have “weaker” surround suppression than the young cells. Third, the mean minimum firing rates did not change between the two conditions, also leading to an interpretation of similar effects of surround in both conditions and cell types. Last, and importantly, the tuning widths in the surround-present condition for old and young cells were globally similar to the no-surround condition, also supporting similar surround effects in both types of neurons.

From this physiological data re-analyses, we found that old cells exhibited higher minimum firing rates but no significant differences in tuning widths when compared to young cells. This confirms our behavioral and modeling results in addition to comforting the previous psychophysical studies.

## Discussion

Neurophysiological studies on orientation processing in the primary visual cortex of animals have proposed that inhibitory function generally declines, and thus healthy older humans might have worse early perceptual abilities than younger persons. However, various psychophysical reports have shown no changes in perceptual capabilities as orientation processing or even opposite effects, where interpretation leads to stronger inhibition. Firstly, we investigated these issues through a combination of psychophysical and modeling approaches. It was found that the amount of center-surround tilt repulsion, attributed to lateral inhibitory interactions and local orientation tuning widths, in the older group was higher compared to younger observers, and that contrast sensitivity of the elder was also globally lower than younger adults. We found a common explanation of all these phenomena in a single model of V1 that dissociated lateral and local inhibitory effects on the perceptual outcome of tilt perception. Our behavioral results and computational modeling provide important evidence that low-level processing deficits in the visual system of elders could be attributed to stronger lateral inhibition. Additionally, the modeling predicted that orientation tuning width in V1 globally should not change with aging. Therefore, secondly, we re-analyzed previous physiological data published by our laboratory and found that the neuronal data followed that prediction. That is, the minimum firing rate of the tuning curve had a strong increase, while orientation tuning widths seemed globally stable across age. Thus, our psychophysical, modeling and re-analysis of neurophysiological results consistently revealed a detail picture of age-related changes in orientation processing, which solved the contradiction between neurophysiological and behavioral reports, and uncovered a differential age-related effects on local inhibition.

Lateral inhibition between neural mechanisms tuned to different orientations in V1 was proposed as an explanation of the repulsive tilt illusion already more than forty years ago(Blakemor.C et al., 1970; Blakemore et al., 1973; Georgeson, 1973). An overall increase in illusory bias of tilt observed in the elder group suggested an increase in lateral inhibition within V1 during aging of early visual function, which was confirmed by the model results, which revealed an increase in lateral inhibition. Our results are in support of recent reports that demonstrated an overall increase in perceptual surround suppression in the older adults when compared to younger observers(Karas and McKendrick, 2015). We reproduced these previous findings with our own stimuli by conducting a center-surround contrast discrimination task for all participants in our study, which correlated to the tilt biases of the subjects. Thus, the similarity in results and interpretation of these two indexes of tilt repulsion and contrast suppression support the idea that they allow to measure a common neural mechanism, and both could be interpreted as stronger lateral inhibition, which should be dissociated from the standard interpretation of reduced local inhibition.

The model, based on the tilt repulsion data, also predicted that the orientation tuning widths of neurons (corresponding to local inhibition (Somers et al., 1995; Shapley et al., 2003)) were similar between the elder and the younger population. These findings account for psychophysical results that sensitivities for orientation processing of the elder are not systematically and substantially changed when compared to younger adults (Betts et al., 2007; Delahunt et al., 2008; Govenlock et al., 2009). In one study (Govenlock et al., 2009), using a typical orientation masking paradigm that indirectly relates to the underlying orientation tuning, the authors did not find differences in tuning widths between groups of young and old persons, as in our results. Although our raw orientation thresholds were different between the two aged groups, these raw results should not be misinterpreted as evidence of differences, due to no adjustment of these values to the individual sensitivity. Indeed, the previous reports (Delahunt et al., 2008) showed that orientation discrimination thresholds are similar across age groups once they are adjusted for inter-subject variability of detection sensitivity, and thus age, which we could not perform due to lack of measurement in our experimental design. Instead, our methods allowed us to infer the underlying theoretical population tuning widths through the modeling of the tilt repulsion effect (Gilbert and Wiesel, 1990b; Tzvetanov, 2012) and thus bypass the long measurement procedures of the previous studies.

The computational prediction that orientation tuning width of neurons were unchanged in the elder when compared to younger populations was confirmed by re-analyzing our previous neurophysiological data collected in young and senescent macaques. These neurophysiological data was collected from classical receptive field measures, which related to local inhibition in our model. The results showed an increased minimum firing rate of the tuning curve and spontaneous activities, which we consider here as neural noise, and maintained orientation tuning widths with aging. Additionally, the agreement of results with a second data set leads to a consistent and deeper understanding of neuronal tuning changes and aging effects onto orientation processing. These outcomes suggest that the two levels of local inhibition, an untuned inhibition affecting neural noise and a tuned inhibition influencing orientation tuning width (Somers et al., 1995; Shapley et al., 2003), were changed by aging differentially, with a weaker untuned local inhibition but a maintained tuned inhibition. These findings revealed a more complex picture of age-related effects in the inhibitory circuits than previously thought.

At the neurophysiological level, previous studies from our laboratory reported an age-related increase in spontaneous neuronal activity in macaque area V1(Leventhal et al., 2003; Wang et al., 2005; Zhang et al., 2008), cat area A17(Hua et al., 2006; Wang et al., 2014) and rat area A17(Wang et al., 2006). One mechanism hypothesized to underlie such changes was a reduction of neuronal/local inhibition due to diminished GABAergic neurotransmission (Leventhal et al., 2003; Wang et al., 2005; Zhang et al., 2008). Since weaker orientation tuned inhibition should strongly broaden the tuning widths (Somers et al., 1995; Shapley et al., 2003), which we did not observe, we propose that a strong weakening of untuned inhibition with aging provides an interpretation of the previously observed changes, while the tuned inhibition remains intact.

However, lateral inhibition, which involves long-range interactions among neurons, seems to increase with aging. The weak GABA mediated local inhibition might result in much higher background firing rates. Hence, a speculative possibility is that an increase in the lateral inhibition is created to counterbalance the decline in local inhibition. However, this “chain of effects” hypothesis needs further research.

In view of previous reports showing a decrease in surround suppression in older adults during motion discrimination task(Betts et al., 2005), the finding of elder adults displaying a stronger center-surround inhibition in tilt illusion and contrast suppression tasks supports the proposal that there is an inhibitory process in the “static” domain of V1 that is different from the mechanism of dynamic motion integration in MT. In primate motion processing, center-surround receptive fields are particularly common in MT(Born and Bradley, 2005; Anton-Erxleben et al., 2009) except the input layer IV(Raiguel et al., 1995), which is one of the evidences indicating that surround inhibition observed in area MT might not be inherited from its feed-forward inputs. Using the same dynamic direction discrimination task originally reported by Tadin and colleagues(Tadin et al., 2003), it was found(Tadin, 2015) that there seem to be a direct relationship between area MT/V5 and spatial suppression. Additionally, they found that when V1 processing was disrupted through TMS pulses performance did not change. On the other hand, lateral inhibition in V1 between neural mechanisms tuned to different orientations was suggested underlie the repulsive tilt illusion (Blakemor. C et al., 1970; Georgeson, 1973). Thus, these two psychophysical measures seem to estimate different aspects of cortical function and seem to reflect independent neuronal mechanisms.

In conclusion, our behavioral, computational and physiological findings provide a new perspective on aging of the visual system. We unveiled two different types of aging-related changes at the primary visual cortex, increase in lateral inhibition and unchanged local orientation coding capacities, but associated to higher neuronal noise. This suggested that local and lateral inhibition were differently affected by aging, and explained disparate results among previous behavioral and neurophysiological studies.

## Acknowledgements

This work was supported by the National Natural Science Foundation of China (31230032 to Y.Z.), General Financial Grant from the China Postdoctoral Science Foundation (2015M571940 to Z.W.) and the Fundamental Research Funds for the Central Universities (to T.T.).

The authors declare no competing financial interests.

